# Delineating genotypes and phenotypes of individual cells from long-read single cell transcriptomes

**DOI:** 10.1101/2023.01.24.525264

**Authors:** Cheng-Kai Shiau, Lina Lu, Rachel Kieser, Kazutaka Fukumura, Timothy Pan, Hsiao-Yun Lin, Jie Yang, Eric L. Tong, GaHyun Lee, Yuanqing Yan, Jason T. Huse, Ruli Gao

## Abstract

Single-cell nanopore sequencing of full-length mRNAs (scNanoRNAseq) is transforming singlecell multi-omics studies. However, challenges include computational complexity and dependence on short-read curation. To address this, we developed a comprehensive toolkit, scNanoGPS to calculate same-cell genotypes-phenotypes without short-read guidance. We applied scNanoGPS onto 23,587 long-read transcriptomes from 4 tumors and 2 cell lines. Standalone, scNanoGPS accurately deconvoluted error-prone long-reads into single-cells and single-molecules. Further, scNanoGPS simultaneously accessed both phenotypes (expressions/isoforms) and genotypes (mutations) of individual cells. Our analyses revealed that tumor and stroma/immune cells often expressed significantly distinct combinations of isoforms (DCIs). In a kidney tumor, we identified 924 genes with DCIs involved in cell-type-specific functions such as *PDE10A* in tumor cells and *CCL3* in lymphocytes. Moreover, transcriptome-wide mutation analyses identified many cell-type-specific mutations including *VEGFA* mutations in tumor cells and *HLA-A* mutations in immune cells, highlighting critical roles of different populations in tumors. Together, scNanoGPS facilitates applications of single-cell long-read sequencing.

## Introduction

Human tumors represent complex ecological systems of diverse cell types with dynamic genetic evolution and phenotypic remodeling^1, 2, 3^. However, there is a lack of robust methods for tracking both genotypes (e.g., mutations) and phenotypes (e.g., gene expressions, isoforms) of individual cells to precisely trace the cellular and molecular dynamics during tumor evolution and treatment response. Long-read single cell sequencing of full-length RNAs is transforming single cell multi-omics analysis through direct measurement of nucleotide sequences of whole gene bodies without algorithmic reconstructions^4, 5, 6, 7^. Nowadays, high throughput long-read single cell sequencing methods take advantage of droplet barcoding systems (commonly, 10X Genomics Chromium system) to barcode full-length cDNAs of single cells and sequence them on ultra-high yield third-generation sequencing (TGS) platforms^5, 6, 7, 8^. Of note, the Oxford Nanopore Technology (ONT) platform, PromethION can yield ~100 million reads per flow cell, providing adequate coverages of thousands of single cell transcriptomes. The PacBio system, Sequel II system can yield ~8 million high fidelity reads, which can measure hundreds of single cell transcriptomes. However, the broad applications of this powerful technology are computationally challenged due to the complexity of calculating same-cell multi-omics from these long-read data. Moreover, due to higher error rates in cell barcodes (CBs) and unique molecule identifiers (UMIs) comparing to next-generation sequencing (NGS), current methods rely on generating paralleled NGS short-read data to guide the deconvolution of CBs and UMIs^5, 8^, which can drastically increase experimental costs and computational complexity and often results in partial usage of data. Recently, ONT released a method, called Sockeye^9^ to perform independent deconvolution of raw reads without the usage of short-reads. However, this method has limited power in thresholding true cells through merging CBs with errors, and needs separate pipelines to calculate single cell multi-omics profiles^10^. Therefore, a robust computational tool for analyzing high throughput single cell long-read data is still missing.

In this study, we developed a computational tool, scNanoGPS (<u>**s**</u>ingle <u>**c**</u>ell <u>**Nano**</u>pore sequencing analysis of <u>**G**</u>enotypes and <u>**P**</u>henotypes <u>**S**</u>imultaneously), to perform independent deconvolution of error-prone long-reads into single-cells and single-molecules and calculate both genotypes and phenotypes in individual cells from high throughput single cell nanopore RNA sequencing (scNanoRNAseq) data. In concert with this tool, we removed all NGS sequencing steps and increased throughput to 3,000-6,000 transcriptomes per flow cell (PromethION). We demonstrated the accuracy and robustness of scNanoGPS and identified cell-type-specific isoforms and mutations in addition to gene expression profiles, enabling synchronous cell-lineage (genotype) and cell-fate (phenotype) tracing to investigate human tumors.

## Results

### Computational workflow of scNanoGPS in analyzing scNanoRNAseq data

The high throughput scNanoRNAseq data are generated through 2 major steps: i) barcoding full-length cDNAs of single cells/nuclei using high throughput droplet barcoding system (10X Genomics), ii) performing high throughput long-read sequencing using PromethION (Oxford Nanopore Technologies) (**Fig. 1a, Online Methods**). Accordingly, the computational workflow of scNanoGPS begins with quality control and scanning reads that have expected patterns of adaptor sequences, i.e., TruSeq R1 adaptor on 5’-ends and TSO adaptor on 3’-ends of raw reads (**Fig. 1a-b, Fig. S2, Online Methods**). Next, we developed an algorithm called <u>**i**</u>ntegrated <u>**C**</u>rude <u>**A**</u>nchoring and <u>**R**</u>efinery <u>**L**</u>ocal <u>**O**</u>ptimization (iCARLO) to detect true cell barcodes (CBs). In this algorithm, a raw list of CBs is determined by searching for a crude anchoring point through thresholding partial derivatives of the supporting reads of individual CBs. The threshold is then extended from crude anchoring point by an empirical percentage (~10%) to rescue true CBs that have much less reads than others in the same experiment. Next, the CBs within two Levenshtein Distances (LDs) are curated and merged to rescue mis-assigned reads due to errors in CB sequences. CBs with less than 300 genes are filtered out by default. iCARLO is implemented in the “Assigner” function, which outputs a list of true CBs and then deconvolutes all qualified reads into single cell FASTQ files.

**Figure 1.**
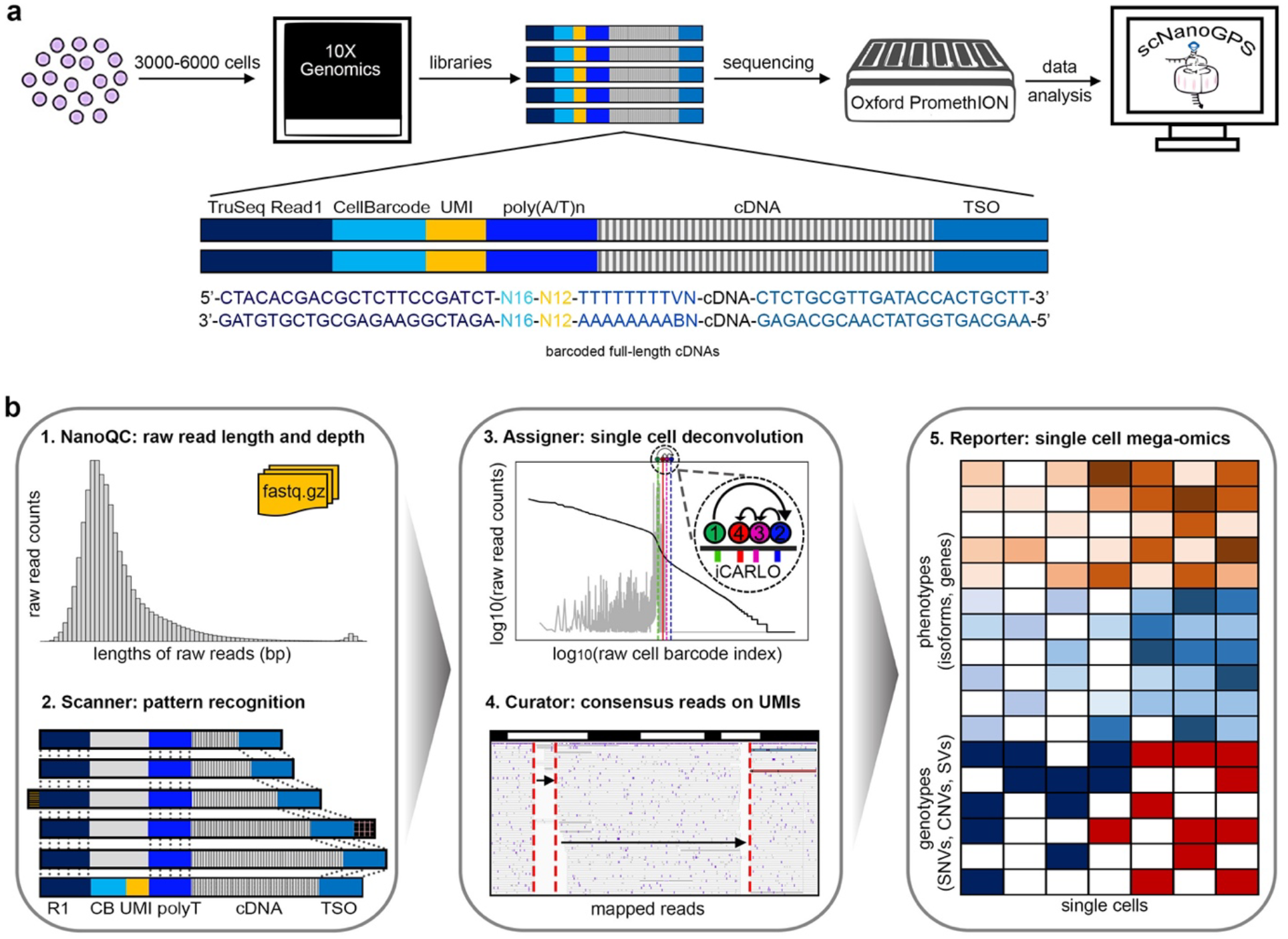
Schematic diagrams of scNanoRNAseq and scNanoGPS workflows. **a**, Experimental workflow and library structure of scNanoRNAseq. **b**, Computational workflow of scNanoGPS.

To identify true UMIs to accurately measure gene expression levels, scNanoGPS first maps single cell reads against reference genome GRCh38^11^ using MiniMap2^12^ (**Fig. 1b**). Reads that map to the same genomic regions (within <5bp) are grouped into read clusters to achieve batch computing. All reads belonging to same read-clusters and having UMIs within two LDs are considered to be amplified from same RNA molecules, thus their corresponding UMIs are curated to be identical. To overcome sequencing errors in single molecules, reads with identical UMIs are collapsed into consensus sequences using SPOA^13^ for better detection of point mutations in gene bodies. At the end, scNanoGPS re-maps single cell consensus reads to generate single cell consensus BAM files and calculates the gene expression, isoform expression and point mutation profiles of individual cells using long-read specific methods^14, 15, 16^. Of note, single cell transcriptome-wide mutation calling results are curated using several criteria to remove random errors (**Online Methods**). Furthermore, single cell copy numbers are calculated using our previously published algorithm CopyKAT^17^. These data can then be used to detect cell-type-specific isoforms and mutations as well as gene expressions to demonstrate the applications of scNanoGPS in investigating the underlying mechanisms of human tumors.

In summary, scNanoGPS independently deconvolutes high throughput scNanoRNAseq data into single-cells and single-molecules without short-read curation and computes same-cell multi-omics profiles for thousands of single cells in parallel.

### Performance of scNanoGPS in processing high throughput scNanoRNAseq data

To test the technical feasibility, we applied scNanoGPS to process the scNanoRNAseq data of two cancer cell lines, A375 and H2030. Strikingly, PromethION yielded 67.4 million reads of the A375 library on one flow cell and 105.3 million reads of the H2030 library on another flow cell. This resulted in averaged depths of 15,944 reads per cell in A375 and 21,710 reads per cell in H2030, which were close to NGS scRNAseq depths. The median read lengths of both datasets were around 900bp, consistent with the traces of pre-sequencing cDNAs (**Fig. 2a, Fig. S1a-b**). The scNanoGPS results showed that most reads (A375: 86.80%, H2030: 87.31%) had the expected pattern of adaptor sequences. In total, we detected 3,649 and 4,212 cells with averaged coverages of 2,688 and 3,553 genes per cell, respectively. To benchmark scNanoGPS performance, we generated standard NGS 3’-scRNAseq (10X Genomics) data from the same cDNA pools as previously described^5, 6^. Comparison analysis showed that scNanoGPS achieved high concordance (92%) with the standard 10X Genomics data in detecting true CBs (**Fig. 2b**), with minor dis-concordance in thresholding low-quality cells that could be mitigated by secondary analyses (**Fig. S3a-b, Online Methods**).

**Figure 2.**
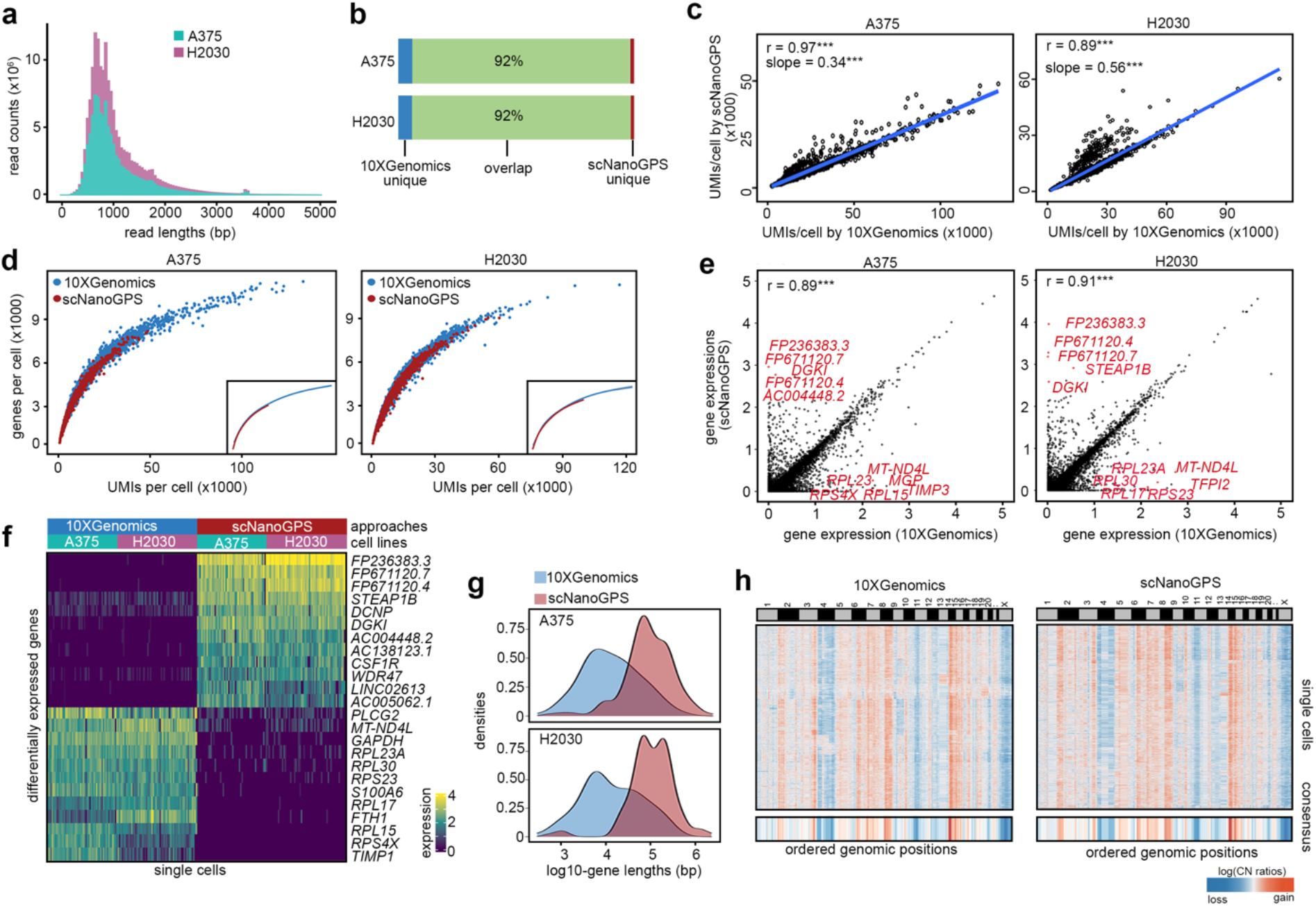
Performance of scNanoGPS in processing scNanoRNAseq data. **a**, Length distribution of raw reads of two cancer cell lines. **b**, Intersection of true cell barcodes detected by scNanoGPS and standard NGS approaches. **c**, Pair-wised scatter plots of UMI counts per cell detected by two approaches. **d**, Pair-wised scatter plots of gene detection versus UMI counts of single cells. **e**, Single cell gene expression levels calculated by two approaches. ***, Pearson correlation P-value <0.001. **f**, Heatmap of genes with significantly different expression levels in two approaches. **g**, Density plots of length of genes with significantly different expression levels in two approaches. **h**, Heatmap of single cell copy number profiles of H2030 calculated by CopyKAT from matched single cell transcriptomes of two approaches.

Next, we compared the number of UMIs detected by scNanoGPS to standard 10X Genomics data. Our results revealed significantly high correlations (A375: Pearson’s r=0.97; H2030: Pearson’s r=0.89) (**Fig. 2c)**. The gene detection rates per UMI were similar as well, although the numbers of UMIs per cell were fewer in long-read data compared to 10X Genomics data (A375: coef=34%; H2030: coef=56%) due to lower sequencing depths (**Fig. 2c-d**).

Further, we compared single cell transcriptomes computed by scNanoGPS to standard 10X Genomics 3’-scRNAseq data. Our results again showed that scNanoGPS achieved high concordance (A375: Pearson’s r=0.89; H2030: Pearson’s r=0.91) in measuring gene expression levels compared to standard 10X Genomics data (**Fig. 2e)**. There were only small fractions (A375: 0.92%; H2030: 0.79%) of detected genes showing significantly different expression levels (FDR-adjusted P-values < 0.05, |log_2_(Fold Changes)| ≥ 1) between the two approaches (**Fig. 2e**). Of note, 10X Genomics 3’-scRNAseq showed a higher chance of detecting ribosomal genes, whereas scNanoGPS detected more long noncoding RNA (lncRNA) genes and pseudogenes (**Fig. 2e-f**). The lengths of scNanoGPS enriched genes were significantly longer than NGS enriched genes (**Fig. 2g.** A375: P-value=8.16×10^-10^; H2030: P-value=1.96×10^-9^), consistent with our expectations.

To investigate whether long-read single cell transcriptome data could serve as a data source for inferring genome-wide DNA copy number alterations (CNAs), we ran CopyKAT^17^ on H2030 which was known to have aneuploidy^16^. As expected, our results showed that H2030 had genome-wide amplifications on Chr2, 3q, 5, 6q, 7, 8q, 14, 15p and deletions on Chr 4, 11p (**Fig. 2h**). The averaged pair-wised Pearson’s correlation between the two approaches reached up to 94%, confirming the feasibility of inferring CNAs using scNanoRNAseq data.

To evaluate whether scNanoGPS can robustly detect major cell types in human tumors, we performed scNanoRNAseq on 4 frozen tumors collected from Renal Cell Carcinoma (RCC1, RCC2) and Melanoma (MEL1, MEL2) patients. We performed unbiased clustering of all cells within each tumor (**Fig. 3a**) and annotated the aneuploid clusters as tumor cell clusters based on CopyKAT results. The tumor cell annotation was validated by the overexpression levels of known cancer genes (**Fig. S4a-b**). The non-tumor cell type clusters were annotated using known celltype markers (**Fig. S4c**). For fair comparisons, we conducted the same clustering and annotations on paralleled 10X Genomics data (**Fig. 3b**). Our results confirmed high concordance between the two approaches (**Fig. 3c**), except that scNanoGPS rescued more lymphocytes that expressed far fewer genes than other cells in the same experiment.

**Figure 3.**
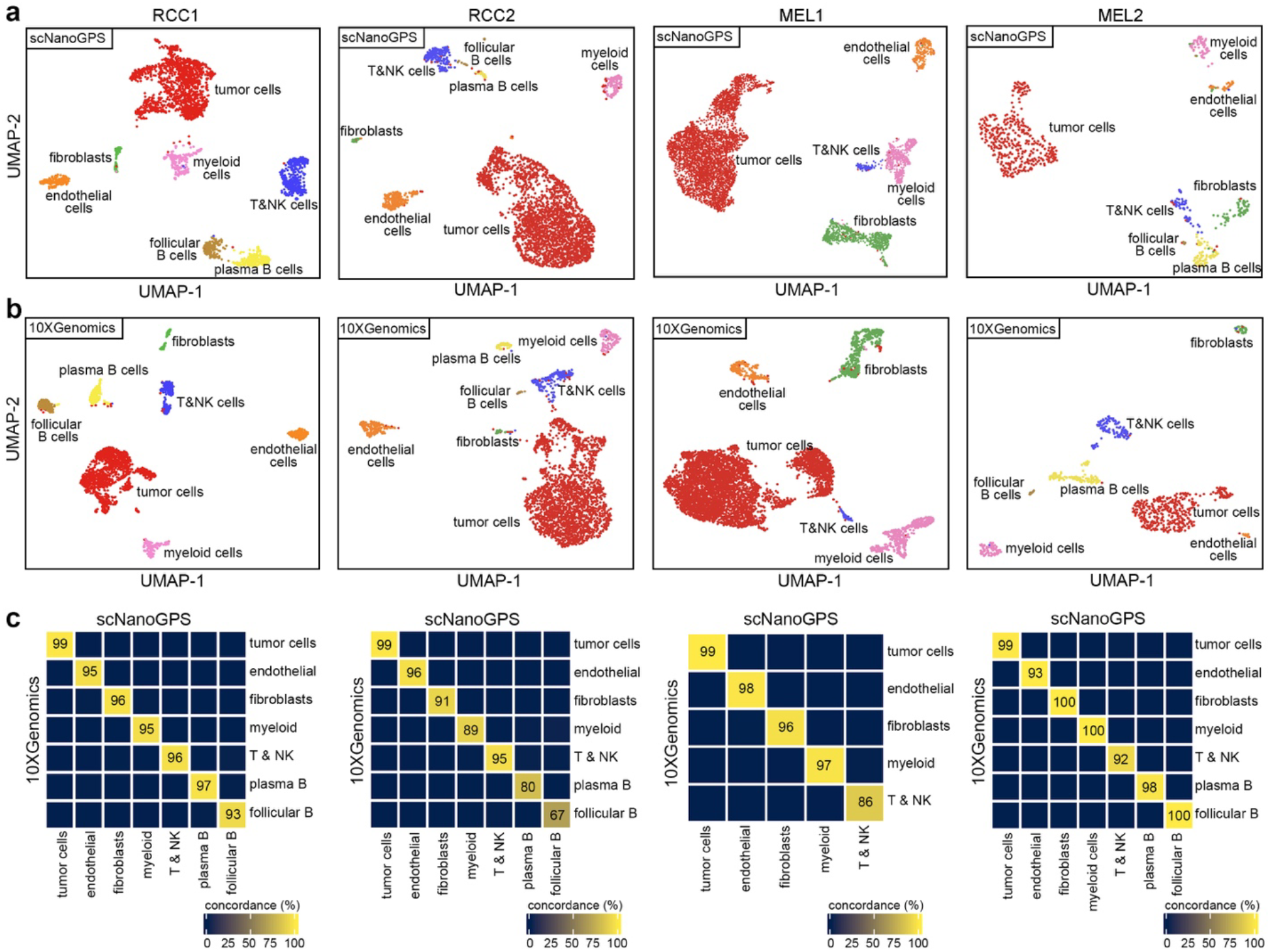
Performance of scNanoGPS in dissecting cell types in the tumor microenvironment. **a**, UMAP projection of major cell types of 4 frozen tumors using data processed by scNanoGPS. **b**, UMAP projection of major cell types of 4 frozen tumors using NGS data. **c**, Concordance of cell typing results between scNanoGPS and NGS approaches. Only cells detected in both approaches were used for calculation.

To summarize, our analyses showed that scNanoGPS reliably deconvoluted long-reads into single-cells and single-molecules to detect single cell transcriptomes and dissect the tumor microenvironment (TME) from scNanoRNAseq data.

### scNanoGPS enables detection of cell-type-specific splicing profiles in human tumors

To access the robustness of scNanoGPS in discovering splicing isoforms of different cell types in the TME, we performed detailed isoform analysis of a frozen kidney tumor (RCC1). On average, we detected 6 transcripts per gene by referring to all known transcripts in GENCODE (*v*32)^18^. Our data showed that each cell type tended to express a different combination of multiple isoforms, consistent with a recent study that reported isoform specificity in mouse cortex^19^. To identify celltype-specific preference of isoforms, we compared the relative compositions of different isoforms of each gene among all 7 major cell types (**Online Methods**). Our analysis identified 1,014 genes that preferably expressed significantly different combinations of isoforms (DCIs, Chi-sq test P-values < 0.05 and |Prevalence Differences| ≥ 10%) among all cell types, including 499 DCI genes in tumor cells, 122 in endothelial cells, 137 in fibroblasts, 90 in myeloid cells, 38 in T & NK cells, 90 in plasma B cells and 38 in follicular B cells (**Fig. 4a-c**). Of note, we detected 2-4 times more genes with DCIs in tumor cells compared to immune and stromal cell types. The top-ranked tumor-cell-specific DCI genes were *PDE10A* and *NR4A2* involved in cAMP pathways. Additionally, we observed that tumor-cell-preferred isoforms had slightly more exons (**Fig. 4d**, paired T-test P-value=0.01), particularly in genes with more than 20 exons such as *NBPF10, VPS13C*, and *NBPF14* (**Fig. 4e**). Gene Ontology (GO) analysis showed that cell-type-specific DCI genes were commonly enriched in pathways related to cell-type-specific functions, such as cAMP pathway and resistance associated glucuronidation pathway in tumor cells, interferon-alpha production pathways in myeloid cells, lymphocyte proliferation pathway in lymphocytes, and immunoglobin-mediated immune responses in B cells (**Fig. 4f**).

**Figure 4.**
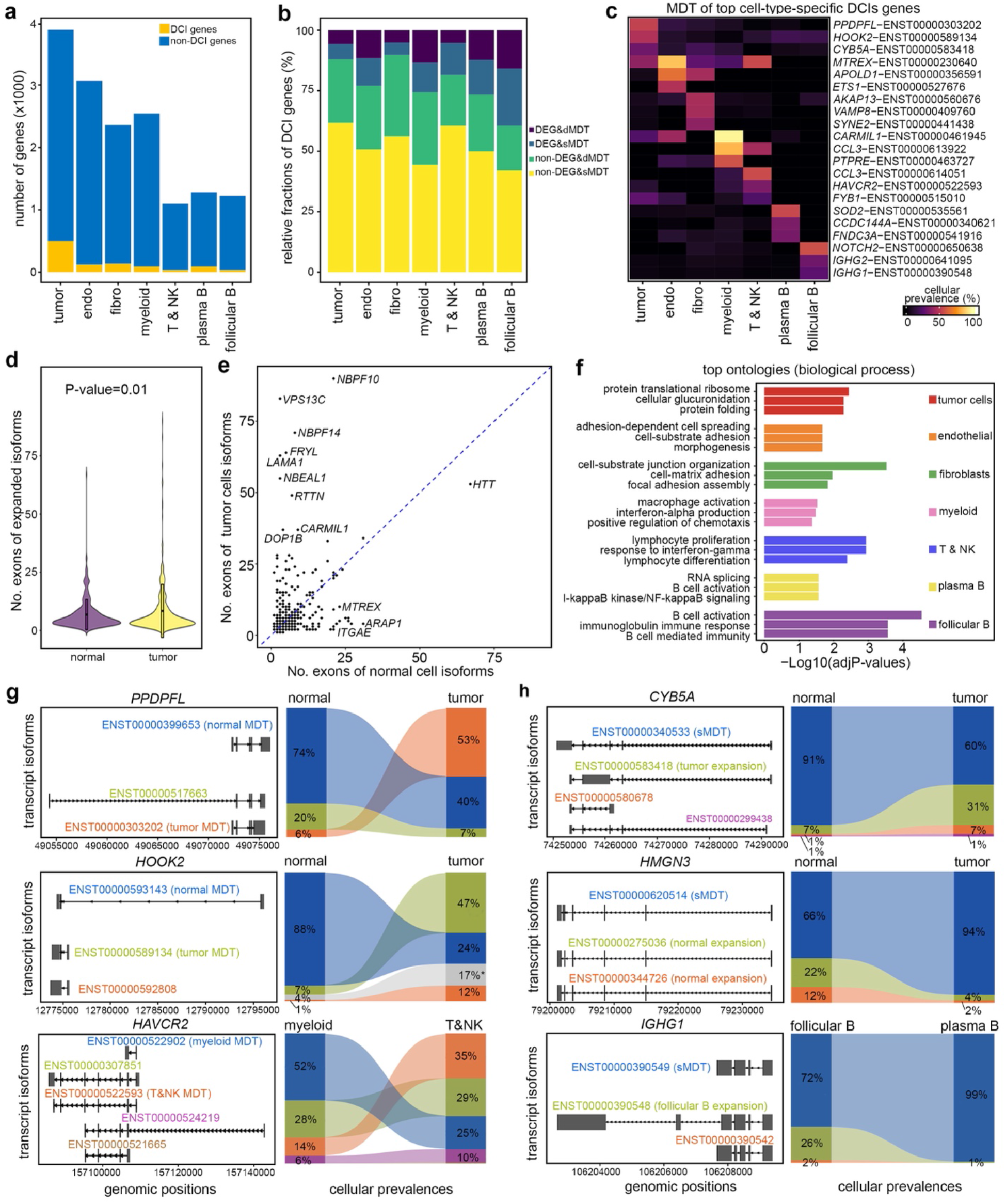
Profiling cell-type-specific isoforms in the tumor microenvironment. **a**, Numbers of genes with DCIs in 7 cell types of a kidney tumor. **b**, Stratification of genes with DCIs based on status of gene expression levels and most dominant transcripts (MDTs). **c**, Heatmap of the cellular frequencies of top cell-type-specific MDTs. **d**, Numbers of exons of tumor and normal cell preferred isoforms. **e**, Pair-wised scatter plot of the numbers of exons of expressed genes in tumor and normal cells. **f**, Top gene ontologies of cell-type-specific DCI genes. **g**, Examples of cellular frequencies of isoforms of genes expressing different MDTs in different cell types. **h**, Examples of cellular frequencies of isoforms of genes expressing same MDTs in different cell types.

Notably, we observed that a large portion of cell-type-specific DCI genes were not detected as differentially expressed genes (DEGs) in all cell types (**Fig. 4b**). Tumor-cell-specific DCI genes preferably expressed different most dominant transcripts (MDTs)^20^ regardless of their overall gene expression levels. For instance, the proliferation gene *PPRPFL* expressed MDTs ENST00000303202 in tumor cells and ENST00000399653 in normal cells, although the overall expression levels of this gene in the two cell types were not significantly different (**Fig. S5a-b**, **Fig.4g**). On the other hand, the organelle hook protein coding gene *HOOK2* preferably expressed ENST00000589134 in tumor cells and ENST00000593143 in normal cells and had significantly higher expression levels in tumor cells compared to normal cells (**Fig. 4g, Fig. S5b**). In addition, we observed a small fraction of cell-type-specific DCI genes that expressed same MDTs, but their cellular fractions were different between tumor and normal cells. One example was *CYB5A* which expressed isoform ENST00000340533 in 60% of tumor cells but 91% of normal cells. Another example was *HMGN3* which expressed isoform ENST00000620514 in 94% of tumor cells but 66% of normal cells (**Fig. 4h, Fig. S5c**).

Interestingly, we also observed isoform preference in immune and stromal cell types, although fewer genes were involved compared to tumor cells. For instance, myeloid and T cells both expressed *HAVCR2*, yet the MDTs of this gene were distinct in the two immune cell types (**Fig. 4g, Fig. S5b**). In contrast, the B cell specific gene, *IGHG1* expressed the same MDT (ENST00000390549) in both follicular and plasma B cells, but the cellular prevalence of this MDT were significantly different in the two B cell subtypes (72% in follicular B cells, 99% in plasma B cells) (**Fig. 4h**), indicating its relevance to sub-cell-type specific functions.

In summary, we demonstrated the usages of scNanoGPS in studying cell-type-specific splicing isoforms in tumors. Our results showed that a larger portion of genes utilized different MDTs in both tumor and immune cell types. Genes expressing same MDTs may have distinct cellular prevalence in different cell types regardless of overall gene expression levels.

### Transcriptome-wide mutations of different cell types in the tumor microenvironment

To accurately detect transcriptome-wide mutations in single cells, we built consensus sequences of single molecules and required at least 2 variant supporting reads. In addition, we filtered out mutations that were detected in less than 1% of cells over all cells or less than 5% within individual cell types (**Online Methods**), which removed most random errors. In total, we detected 6,390 mutations from 3,470 single nuclei transcriptomes of a frozen kidney tumor (RCC1). Among all mutations, 17.9% were in exonic regions, while others were spreading across non-coding regions (35.8% intronic, 28.8% intergenic, 3.7% 5’UTR and 13.8% 3’UTR) (**Fig. 5a**). Of note, we observed 53.3% exon mutations were nonsynonymous. Our results showed that transition mutation types were more frequent (each 13-15%) than transversion mutation types (each 4-6%) (**Fig. 5b**), consistent with a previous study^21^. The potential RNA editing (C>U) events^22^ were not distinguishable from C>T transition (15.3%). Our data showed that these transcriptome-wide mutations were distributed across all chromosomes except for Y-chromosome that expressed far fewer genes (**Fig. 5c**). Interestingly, we identified 4 shared mutation hotspots on Chr 2, 6, 14 and 22 in all 7 cell types, which resulted in frequent mutations on microRNA, HLA and lncRNA genes (**Fig. S6a-b**).

**Figure 5.**
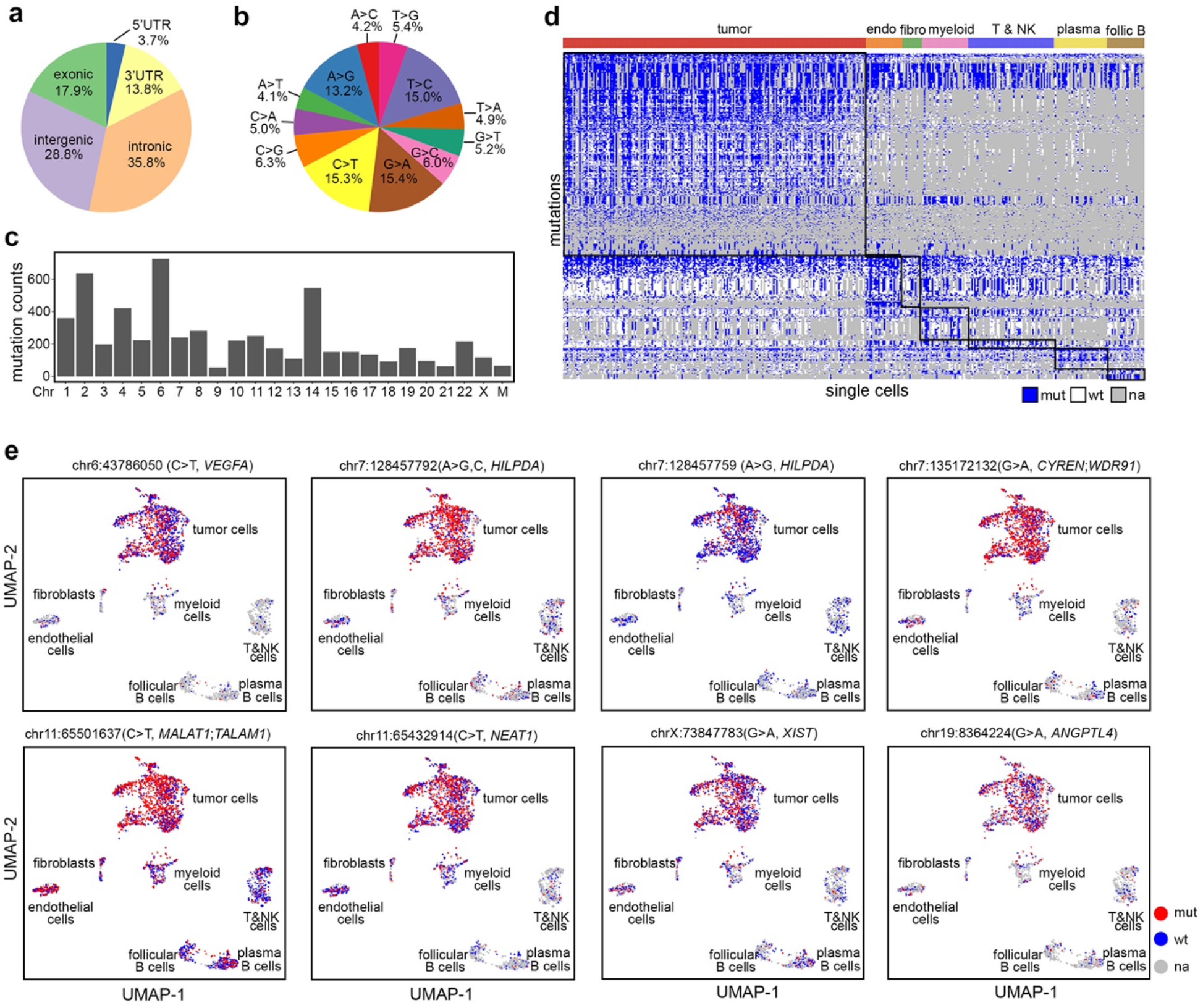
Transcriptome-wide mutation profiling of different cell types in the tumor microenvironment. **a**, Pie chart of relative fractions of all mutations in different gene regions. **b**, Pie chart of mutation types of all mutations. **c**, Number of mutations in different chromosomes. **d**, Heatmap of cell-type-specific deMuts in all cells. **e**, UMAP projections of single cells labeled with 8 examples of tumor-cell-specific deMuts.

Our data showed that the mutation detection rates were between 15-25% among all cells (**Fig. S7a**), consistent with single cell transcriptome-wide gene detection rates^3, 23, 24, 25^. Among all cells that had detectable gene expression, the frequencies of cells expressing mutants were strikingly high (above 75%) in all cell types (**Fig. S7b**). Of note, we rarely observed mutations that were exclusively detected in one cell type. Instead, they were expanded at varying cellular frequencies depending on cell type. We compared the cellular frequencies of mutations among different cell types to identify mutations that were differentially expanded (deMuts). Our results showed that tumor cells had the largest number (N=609) of deMuts, followed by myeloid cells (N=99), plasma B cells (N=63), endothelial cells (N=57), follicular B (N=29), lymphocytes (N=26) and fibroblasts (N=1) (**Fig. 5d)**. Tumor cell specific deMuts included many COSMIC genes, such as *VEGFA*, *NEAT1*, *MALAT1*, *HILPDA* etc. (**Fig. 5e**). Further, we detected several genes that were both mutated and differentially spliced in the same cell types, such as *HMGN3* and *UBE2G2* in tumor cells, *CD74* and *IFI30* in myeloid cells, and *RPS2* in endothelial cells indicating their important roles in tumorigenicity. Additionally, we observed several cell-type-specific deMuts involved in the spliceosome (GSEA: KEGG pathway) such as *SRSF3* and *HNRNPC* mutants in tumor cells and *DDX5* mutant in endothelial cells.

In summary, we demonstrated that scNanoGPS can robustly detect cell-type and cellstate specific mutations. Our data highlighted the importance of identifying population-specific mutations to understand their functional roles in cancer progression.

## Discussion

In this study, we developed a computational tool called scNanoGPS to facilitate high throughput single cell long-read sequencing analysis of human tumors. scNanoGPS addresses the major computational challenges of an emerging powerful technology. It achieves independent deconvolution of raw data without the guidance of short-reads and calculates genotypes-phenotypes of thousands of individual cells with accuracy and robustness.

Two previous methods called Sicelore^5^ and scNapBar^8^ were developed to deconvolute raw reads, however, both relied on the guidance of paralleled NGS data. An experimental method called scCOLOR-seq^7^ was developed to reduce error rates in barcodes by designing bi-nucleotide repeats in barcode sequences^7^, but it still relied on prior knowledge of true barcodes to curate errors. Moreover, scCOLOR-seq required customized synthesis of gel-beads to adapt to the droplet system. The ONT Sockeye^9^ performed independent deconvolution of raw reads without the usages of short-reads by directly merging CBs within certain editing distance, which thus far has limited power in thresholding true cells. Additionally, it needs users to build separate pipelines to calculate isoforms and mutations. In contrast, scNanoGPS achieves independent data deconvolution, corrects errors in gene bodies, and calculates same-cell mutations, isoforms, and gene expressions simultaneously for thousands of cells.

Dysregulation of transcript isoforms plays critical roles in tumor progression^20, 26, 27, 28^. Long-read single cell sequencing technologies enable in-depth annotation of splicing isoforms at single cell levels. scNanoGPS provides a robust computational tool to achieve this goal. Our analysis of a frozen kidney tumor led to paradigm shift findings, i.e., all major cell types in the tumor express a combination of different isoforms instead of one canonical isoform. Another important finding is that tumor cells preferably express different MDTs of tumor suppressors although their overall gene expression levels are not significantly different from normal cells. Our results implied that the discovery of cancer-specific genes using gene expression levels may only be revealing the tip of iceberg of transcriptional diversities in cancer.

scNanoGPS provides a powerful approach for synchronic tracing of cell-lineage and cellfate by measuring both plastic phenotypic markers (genes, isoforms) and stable genetic markers (mutations, copy numbers) of the same cells to study tumor evolution and therapeutic responses. scNanoGPS detects transcriptome-wide point-mutations with accuracy by building consensus sequences of single molecules and performing consensus filtering of cellular prevalence, which removes most false calls due to random sequencing errors. However, our consensus approach does not address errors in calling small indels that represent the major type of Nanopore sequencing errors. We expect that new versions of sequencing chemistry and nucleotide calling algorithms could address this limitation.

In addition to what we have demonstrated in this study, scNanoGPS has broad applications in many other genomic areas, such as measuring single cell gene fusions, tandem repeats, splicing velocities, repetitive genes, or long non-coding genes to investigate diverse mechanisms of human diseases including but not limited to cancer.

## Data and software availability

Raw sequencing data and processed data are submitted to Gene Expression Omnibus (GEO): GSE212945. Software is available at GitHub (https://github.com/gaolabtools/scNanoGPS).

## Acknowledgements

We thank research funding supports to R.G. from Northwestern University and National Heart, Lung, and Blood Institute (NIH 1R01HL160552-01). The nanopore sequencing was carried out by the DNA Technologies and Expression Analysis Core at the UC Davis Genome Center, supported by NIH Shared Instrumentation Grant (1S10OD010786-01). The next generation sequencing was carried out by the ATGC core at University of Texas MD Anderson Cancer Center, supported by the Core grant (CA016672) and NIH Shared Instrument Grant (1S10OD024977-01), and by the Genomic and RNA Profiling Core at Baylor College of Medicine with funding support from an NIH S10 grant (1S10OD023469).

## Author contributions

C.S., L.L., and R.K. contributed equally to this work. C.S. led the development of scNanoGPS toolkit, performed data analysis and wrote the manuscript. L.L. led the development of methods for analyzing cell-type-specific isoforms and mutations, performed data analysis and wrote the manuscript. R.K. led the development of scNanoRNAseq workflow, generated data, reviewed results and wrote manuscript. F.K. cultured cell lines, collected human tissues, and reviewed results. T.P. reviewed results and wrote manuscript. H.L. performed experiments and wrote manuscript. J.Y. performed data analysis and reviewed results. E.T. analyzed data and reviewed results. G.L. performed experiments. Y.Y. conceived concepts and reviewed results. J.H. designed and supervised collection of clinical samples and cancer cell lines, reviewed results, and edited manuscript. R.G. conceived concepts, supervised overall study, designed computational tools and experiments, analyzed data, reviewed results, and wrote manuscript.

## Methods

### Cancer cell line and tumor tissue samples

The two cancer cell lines (A375, H2030) and the four frozen tumors (RCC1, RCC2, MEL1, MEL2) were provided by Dr. Jason Huse at UT MD Anderson Cancer Center. The cell lines are standard commercialized cancer lines. The tumor tissues were collected with consent under IRB approval at UT MD Anderson Cancer Center.

### Preparation of single nucleus suspension

Single nucleus suspensions of two cancer cell lines were prepared by following the 10X Genomics protocol (CG000365 Rev C). Single nucleus suspensions of frozen tumor tissues were prepared according to the method as previously described^24, 29^. Frozen tissue was cut into tiny pieces in 10-cm Petri dish with 500ul-2ml NST-DAPI buffer for 10-15 minutes and filtered through a 40 mm Flowmi into 1.5ml LowBind Eppendorf tube and centrifuged at 4°C 300g for 5 minutes. The resulting nuclei pellet was washed three times with cold Nuclei Wash and Resuspension Buffer. After cell counting, the nuclear suspension was centrifuged again and resuspended in the appropriate volume depending on the nuclei counting results. Preparation of NST-DAPI buffer: Mix 800 ml of NST solution (146 mM NaCl, 10 mM Tris-base (pH 7.8), 1 mM CaCl2, 21 mM MgCl2, 0.05% (wt/vol) BSA and 0.2% (*v/v*) Nonidet P-40) with 200 ml of DAPI solution (106 mM MgCl2 and 10 mg of DAPI). The solution is filter-sterilized and is stored at 4°C in the dark for up to 1 year. Preparation of Nuclei Wash and Resuspension Buffer: 1X PBS with 1.0% BSA and 0.2U/μl RNase Inhibitor.

### Preparation of barcoded full-length cDNAs of single nuclei

Single nucleus suspensions were loaded onto 10X Genomics Chromium Controller (iX) with Chip J to capture 3000-6000 single nuclei. The full-length mRNAs and/or pre-mRNAs were barcoded with cell barcodes (cellBCs) and unique molecular identifiers (UMIs) through cDNA amplification using 10X Genomics protocol. We modified the cDNA amplification protocol by extend the elongation time to 3 minutes to enrich longer molecules as previously described^5^.

The barcoded full-length cDNA transcripts (10ng) were amplified for five cycles with the following two customized primers synthesized by Integrated DNA Technologies (Coralville, IA): 1) 5’-biotinylated TruSeq Read 1 forward primer 5’-/5Biosg/AA AAA CTA CAC GAC GCT CTT CCG ATC T −3’ (25 nM); 2) 3’ partial TSO reverse primer 5’-NNN AAG CAG TGG TAT CAA CGC AGA GTA CAT-3’. The amplified single-cell cDNA transcripts were subjected to 0.8X SPRIselect reagent (Beckman Coulter, CA) clean-up to remove unbound and excess biotinylated primers, where the bound cDNA was eluted off the bead matrix in 45 μL of Qiagen Buffer EB (Qiagen; Valencia, CA). The eluted cDNA was further purified through the binding of the biotinylated template to Dynabeads™ M-270 Streptavidin beads (Invitrogen; Waltham, MA). Prior to the selection of the cDNA template, 15 μL of the Dynabeads™ M-270 Streptavidin beads were washed in 1 mL of 1X SSPE solution (UltraPure™ 20X SSPE Buffer (Invitrogen) freshly prepared with nuclease-free water). A magnetic stand was used to separate the streptavidin beads from the initial wash solution. The streptavidin beads were then washed three times with 15 μL of 1X SSPE with removal from the magnet and resuspension of the beads in a fresh wash solution with each repeat wash. Following the final wash, the streptavidin beads were resuspended in 10 μL of 5X SSPE solution and the biotinylated cDNA template was added to the washed beads. This mixture was placed on a tube rotator at room temperature for 15 minutes. Post incubation on the rotator, with the biotinylated cDNA template bound to the washed streptavidin beads, the sample was placed back on the magnetic stand for separation. The cDNA-bound beads were washed twice with 100 μL 1X SSPE solution and a final wash with 100 μL Buffer EB. The use of the biotinylated forward primer and subsequent purification with the Dynabeads™ M-270 Streptavidin beads allowed for the selective depletion of cDNA missing the terminal poly(A)/poly(T) tail.

The streptavidin beads containing bound biotinylated cDNA were then resuspended in 100 μL of PCR master mix for a secondary amplification for five cycles with regular PCR primers: 1) TruSeq read 1 forward primer 5’-NNN CTA CAC GAC GCT CTT CCG ATC T-3’ and 3’ partial TSO reverse primer 5’-NNN AAG CAG TGG TAT CAA CGC AGA GTA CAT-3’. The PCR amplified product was purified with 0.8X SPRIselect reagent into a final elution of 51 μL in Buffer EB to allow for adequate template/volume for the necessary assessment of quality control (QC) metrics and PromethION library preparation.

The KAPA Biosystems HiFi HotStart PCR Kit (Roche; Basel, Switzerland) was used to prepare all PCR amplification mixtures for Nanopore library preparations. The following PCR conditions were followed for amplification: initial denaturation, 3 mins at 95°C; five cycles of denaturation for 30 secs at 98°C, annealing for 15 secs at 64°C, and extension for 5 mins at 72°C; followed by a final extension for 10 mins at 72°C.

### Nanopore sequencing library preparation for full-length cDNAs

Based on the sample molarity and average cDNA transcript length derived from the QC metrics, the sample input volume was calculated and used to progress into nanopore library preparation. The SQK-LSK110 ligation sequencing kit (Oxford Nanopore) was used to generate PromethION long-read cDNA libraries. The final sequencing was run on PromethION flow cell (*v9.4.2*) with one sample per flow cell by the DNA technology core at UC Davis using R9.4 chemistry. The output data was base-called live during the run using base-caller guppy (*v5.0.12*) in super-accurate base-calling model.

### Generating benchmarking data with paralleled NGS sequencing of fragmentized cDNAs

The same aliquot (25% volume) of barcoded full-length cDNAs were fragmented and subjected to next generation sequencing library preparation by following 10X Genomics Next GEM Single Cell Gene Expression protocol. The final libraries were sequenced on the Illumina Novaseq 6000 sequencer at the NUcore at Northwestern University. The sequencing data were processed using 10X Genomics software CellRanger ARC (*v2.0*)^25^.

### Quality control of scNanoRNAseq data

The raw FASTQ files were processed by the scNanoGPS ‘NanoQC’ function to scan the distribution of raw read lengths, which generated a PNG plot named ‘read_length.png’ and a tab-separated table named ‘read_length.tsv’. The first and last 100 nucleotides of raw reads were extracted for sequencing quality analysis FastQC^30^.

### Scanning long-reads with expected adaptor patterns

The raw FASTQ files were processed by scNanoGPS ‘Scanner’ function to scan the expected adaptor patterns using the parameters (match: 2, mismatch: −3, gap opening: −5, gap extension: −2, sequence identity ≥ 70%) equivalent to NCBI Basic Local Alignment Tool (BLAST) algorithm. Raw reads with insert length less than 200 bp were excluded from scanning. After ‘Scanner’, a compressed file containing parsed raw cell barcodes (CBs) named ‘barcode_list.tsv.gz’ and a filtered read sequence file named ‘processed.fastq.gz’ were generated.

### Deconvolution of long-reads into single cells

The raw reads that passed QC and pattern filtering steps were demultiplexed into single cells using scNanoGPS ‘Assigner’ function. Another input was the parsed barcode lists generated by ‘Scanner’. The true list of CBs was retrieved through an integrated algorithm called iCARLO. This algorithm included 4 steps. First, all CBs were ordered decreasingly by the number of supporting reads. The number of supporting reads and the order index were transformed into log10 scale (**Fig. 1b step 3 Assigner**). The raw list of true CBs was estimated by thresholding the maximal partial derivatives of supporting reads against the rank of CBs. To buffer the changes, we smoothed the partial derivatives within each 0.001 window of log10-scaled CB ranks.

Let *X* be the number of supporting reads of CBs, *i*. be the rank order of all CBs, and *w* be the 0.001 smoothing windows as defined in equation (1).

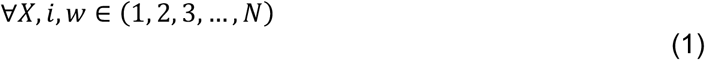

Each window *w* contains a set of CBs, allowing empty, according to their rank order *i*. in log10 scale per 0.001 tick as shown in equation (2).

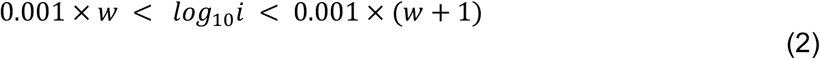

The partial derivatives were calculated for each CB and then smoothed by taking median average of all values within each window *w* as shown in equation (3). The crude anchoring point was defined as a threshold cutoff where a smoothing window *w* had maximal partial derivative. We defined the raw number of CBs (*Cr*) at this crude anchoring point as follows:

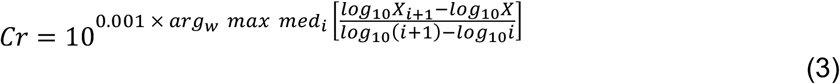

We then extended this crude anchoring point by an empirical percentage (10%) to an external boundary (N= *Cr*_2_) of true CBs to rescue as many CBs as possible. In the third step, we exhaustively computed pair-wised LD distances of all *Cr*_2_ CBs. CBs within 2 LD were merged back one directionally to the CB harboring more supporting reads and obtained a collapsed list of true CBs (N= *Cr*_3_). Lastly, we retrieved the final list of true CBs (N= *Cr*_4_) by removing CBs which cover less than 300 genes. According to this final list of true CBs, the master FASTQ file resulting from ‘Scanner’ was split into single cell FASTQ files stored in a temporal folder holding all the meta files for further usages.

### Curation of sequencing errors in single molecules

Detection of the true list of UMIs and curation of sequencing errors in single molecules was performed by using scNanoGPS ‘Curator’ function. To detect true UMIs, deconvolution of the single cell FASTQ files were first aligned to the reference genome (GRCh38) using MiniMap2^12^ under the splicing mode (-ax splice). Reads mapped to the same genomic regions (coordinates within 5bp) were grouped into batches. Batch calculation was conducted by paralleled computing cores. For the reads, that belong to the same clusters and have UMI within two LDs, were considered as reads of the same molecules. These UMI sequences were curated to be identical. To curate errors in gene bodies, the reads with the same curated UMIs were collapsed into consensus sequences of single molecules using SPOA^13^. Finally, the consensus reads were re-mapped against the reference genome (GRCh38) using MiniMap2^12^ under splicing mode (-ax splice) to generate consensus BAM files for all downstream analyses.

### Calculation of same-cell multi-omics from consensus reads

The consensus BAM files of single cells were used as input to calculate single cell transcriptomes, isoforms and point mutations using scNanoGPS ‘Reporter’ functions. The single cell gene expression profiles were calculated using FeatureCount^14^ from the Subread package^31, 32^ that support long-read gene level counting. The single cell isoforms were calculated using LIQA^15^, a method designed to calculate isoforms from long-read data of spliced mRNAs. We used default parameters (weight of bias correction: 1, maximal distance: 20bp) when running LIQA. The single cell point mutation detection was conducted using a robust long-read variant detection tool, LongShot^16^. We benchmarked the LongShot results in terms of the number of SNVs with different cell prevalence and different number of supporting reads. With the elbow method, we required that each alternative allele should be supported by at least two consensus reads. The resulting VCF files of single cells were merged using BCFtools (*v1.15*)^33^. A final list of mutations of all cells was obtained by consensus filtering, where all variants detected in less than 1% of the cells were considered as random errors and removed from analysis. To differentiate between true wildtypes and missing values, we re-scanned the read depths of all loci in the final list by Samtools (*v1.15*)^33^. Loci with 0/0 genotypes with supporting reads >0 were defined as true wildtypes, otherwise, as missing values if none supporting reads were found. The final mutation loci were annotated using ANNOVAR^34^, which included dbSNP (*v150*)^35^ and COSMIC (*v96*)^36^ as references. The single cell copy numbers were calculated using our previously published method, ‘CopyKAT’ (*v1.0.6*)^17^ with default parameters. In all final output matrices, features/genes were put in rows while cell barcodes were in the columns.

### Single cell gene expression data analysis: QC and defining major cell types

The gene expression matrices of NGS-based 3’-scRNAseq data were processed using CellRanger ARC^25^ (10X Genomics) and sent for downstream analysis using the ‘Seurat’ R package (*v4.1.1*)^37^. In 4 tumor samples, doublets were removed using R package ‘DoubletFinder’ (*v2.0.3*)^38^ with an assumed doublet rates of 0.8% per 1000 cells. Cells with more than 10,000 genes, or more than 100,000 UMIs, considered as doublets, were removed as outliers and suspected doublets. Cells with less than 300 genes were filtered out due to low gene coverages. Furthermore, cells with extremely higher fractions of mitochondrial genes were filtered out using arbitrary outlier cutoffs (5% in RCC1 and RCC2; 20% in MEL1 and MEL2). UMI count matrices were normalized using ‘LogNormalize’ method in ‘NormalizeData’ function and scaled across cells using ‘ScaleData’ function in ‘Seurat’. The top 2000 highly variable genes were selected with ‘FindVariableFeatures’ function based on ‘vst’ method and used for Principal Component Analysis (PCA). Next, we performed PCA and uniform manifold approximation and projection (UMAP) for dimension reduction with the top 30 PCs. ‘FindNeighbors’ function based on the top 30 PCs and ‘FindClusters’ functions were applied to perform unbiased clustering of cells. In all the samples, we defined a cluster of low-quality cells that did not express known cell type markers and had much fewer genes and UMIs compared to other cells in the same experiments. Final clustering analyses were reperformed without these low-quality cells. To identify the tumor cells, we used the UMI count matrix as input to infer chromosomal CNA profiles using R package ‘CopyKAT’ (*v1.0.6*)^17^. Cells with genome-wide CNAs were labeled as tumor cells.

### Analyses of cell-type-specific splicing isoforms

Cell-type-specific splicing isoform analyses started from single cell isoform expression matrices that summarized the expression levels of all known isoforms based on GENCODE (*v32*)^18^ in single cells. For pairwise two group (cell type) comparisons, we filtered out sporadically expressed genes, which were detected in less than 5% cells in both comparison groups. Additionally, genes with only one isoform were excluded from this analysis. To mitigate false discovery driven by dropouts, the isoform expression levels of a given gene were aggregated across all cells within each comparison group. The aggregated counts of expressed molecules of all isoforms of a given genes in both comparison groups were sent for Chi-square tests to test whether the relative composition of different isoforms of a given gene was significantly different in the two comparison groups or not. P-values were adjusted using Benjamin-Hochberg (BH) correction for multiple testing with a 5% false discovery rate. The relative frequency of all isoforms of a given genes and the differences in two comparison groups were also calculated. Finally, the genes expressing significantly different combination of isoforms (DCIs) were defined as having FDR adjusted P-values < 0.05 and at least one isoform had different cellular prevalence in two comparison groups ≥ 10%.

### Detection of cell-type-specific SNVs

To further remove random errors, we filtered out the called positions that were detected in less than 5% cells in each cell type or in comparing group. Next, we generated a count matrix that included the number of wild-type cells (expressed only reference alleles) and the number of mutated cells (expressed variant alleles) of all candidate SNVs in both groups that were under comparison. We performed a Chi-square test to measure whether a candidate SNV had significantly different cellular frequencies. We adjusted P-values using BH correction for multiple testing with a false discovery rate of 5%. This analysis was only performed using data of cells that had read coverages. Cells without read coverages were not included to calculate the cellular frequencies of mutations. We determined the candidate SNVs with FDR adjusted P-value < 0.05 and the differences in cellular frequencies > 0.1 as population specific differentially expanded SNVs (deMuts).

### Summary of statistical methods

We applied Chi-sq tests to compare the relative frequencies of isoforms or mutations in two comparison groups. P-values were adjusted using BH method to adjust for multiple test errors with a false discovery rate of 5%. We applied Pearson’s correlation and P-value to measure the similarity of gene expression profiles and the total counts of UMIs per cell between scNanoGPS and NGS approaches. Paired-two-side t-test was performed to compare the number of exons of different isoforms of same genes between tumor and normal cells. All significance cutoffs used in this study were set at 0.05.

## References

1. Wang Y, et al. Clonal evolution in breast cancer revealed by single nucleus genome sequencing. Nature 512, 155–160 (2014).

2. Kim C, et al. Chemoresistance Evolution in Triple-Negative Breast Cancer Delineated by Single-Cell Sequencing. Cell 173, 879–893 e813 (2018).

3. Patel AP, et al. Single-cell RNA-seq highlights intratumoral heterogeneity in primary glioblastoma. Science 344, 1396–1401 (2014).

4. Fan X, et al. Single-cell RNA-seq analysis of mouse preimplantation embryos by third-generation sequencing. PLoS Biol 18, e3001017 (2020).

5. Lebrigand K, Magnone V, Barbry P, Waldmann R. High throughput error corrected Nanopore single cell transcriptome sequencing. Nat Commun 11, 4025 (2020).

6. Tian L, et al. Comprehensive characterization of single-cell full-length isoforms in human and mouse with long-read sequencing. Genome Biol 22, 310 (2021).

7. Philpott M, et al. Nanopore sequencing of single-cell transcriptomes with scCOLOR-seq. Nat Biotechnol 39, 1517–1520 (2021).

8. Wang Q, Boenigk S, Boehm V, Gehring NH, Altmueller J, Dieterich C. Single cell transcriptome sequencing on the Nanopore platform with ScNapBar. RNA, (2021).

9. Technologies TON. Sockeye: nanopore-only demultiplexing of single-cell reads. (2022).

10. Technology ON. Workflow single-cell. (2022).

11. Schneider VA, et al. Evaluation of GRCh38 and de novo haploid genome assemblies demonstrates the enduring quality of the reference assembly. Genome Research 27, 849–864 (2017).

12. Li H. New strategies to improve minimap2 alignment accuracy. Bioinformatics, (2021).

13. Vaser R, Sovic I, Nagarajan N, Sikic M. Fast and accurate de novo genome assembly from long uncorrected reads. Genome Res 27, 737–746 (2017).

14. Liao Y, Smyth GK, Shi W. featureCounts: an efficient general purpose program for assigning sequence reads to genomic features. Bioinformatics 30, 923–930 (2014).

15. Hu Y, Fang L, Chen X, Zhong JF, Li M, Wang K. LIQA: long-read isoform quantification and analysis. Genome Biol 22, 182 (2021).

16. Edge P, Bansal V. Longshot enables accurate variant calling in diploid genomes from single-molecule long read sequencing. Nat Commun 10, 4660 (2019).

17. Gao R, et al. Delineating copy number and clonal substructure in human tumors from single-cell transcriptomes. Nat Biotechnol 39, 599–608 (2021).

18. Frankish A, et al. GENCODE 2021. Nucleic Acids Research 49, D916–D923 (2021).

19. Booeshaghi AS, et al. Isoform cell-type specificity in the mouse primary motor cortex. Nature 598, 195–199 (2021).

20. Kahraman A, Karakulak T, Szklarczyk D, von Mering C. Pathogenic impact of transcript isoform switching in 1,209 cancer samples covering 27 cancer types using an isoform-specific interaction network. Sci Rep 10, 14453 (2020).

21. Rosenberg MS, Subramanian S, Kumar S. Patterns of transitional mutation biases within and among mammalian genomes. Mol Biol Evol 20, 988–993 (2003).

22. Ramaswami G, et al. Identifying RNA editing sites using RNA sequencing data alone. Nat Methods 10, 128–132 (2013).

23. Yamawaki TM, et al. Systematic comparison of high-throughput single-cell RNA-seq methods for immune cell profiling. BMC Genomics 22, 66 (2021).

24. Gao R, et al. Nanogrid single-nucleus RNA sequencing reveals phenotypic diversity in breast cancer. Nat Commun 8, 228 (2017).

25. Zheng GX, et al. Massively parallel digital transcriptional profiling of single cells. Nat Commun 8, 14049 (2017).

26. Zhang S, et al. A widespread length-dependent splicing dysregulation in cancer. Sci Adv 8, eabn9232 (2022).

27. Ouyang J, et al. The role of alternative splicing in human cancer progression. Am J Cancer Res 11, 4642–4667 (2021).

28. Liu Q, Fang L, Wu C. Alternative Splicing and Isoforms: From Mechanisms to Diseases. Genes (Basel) 13, (2022).

29. Sei E, Bai S, Navin N. Dissociation of Nuclear Suspensions from Human Breast Tissues. (2018).

30. S A. FastQC: a quality control tool for high throughput sequence data.). Available Online (2010).

31. Liao Y, Smyth GK, Shi W. The Subread aligner: fast, accurate and scalable read mapping by seed-and-vote. Nucleic Acids Res 41, e108 (2013).

32. Liao Y, Smyth GK, Shi W. The R package Rsubread is easier, faster, cheaper and better for alignment and quantification of RNA sequencing reads. Nucleic Acids Res 47, e47 (2019).

33. Danecek P, et al. Twelve years of SAMtools and BCFtools. GigaScience 10, (2021).

34. Wang K, Li M, Hakonarson H. ANNOVAR: functional annotation of genetic variants from high-throughput sequencing data. Nucleic Acids Res 38, e164 (2010).

35. Sherry ST. dbSNP: the NCBI database of genetic variation. Nucleic Acids Research 29, 308–311 (2001).

36. Tate JG, et al. COSMIC: the Catalogue Of Somatic Mutations In Cancer. Nucleic Acids Res 47, D941–D947 (2019).

37. Stuart T, et al. Comprehensive Integration of Single-Cell Data. Cell 177, 1888–1902 e1821 (2019).

38. McGinnis CS, Murrow LM, Gartner ZJ. DoubletFinder: Doublet Detection in Single-Cell RNA Sequencing Data Using Artificial Nearest Neighbors. Cell Syst 8, 329–337 e324 (2019).

