## Supplementary Figures for "Delineating genotypes and phenotypes of individual cells from long-read single cell transcriptomes"

*Shiau, Lu, Kieser et al., 2023*

### Table of Contents

**Figure S1 – Size distributions of cDNAs, related to Figure 1-2**

**Figure S2 – Illustration of scNanoGPS methods, related to Figure 1**

**Figure S3 – Identification of low-quality cells, related to Figure 2**

**Figure S4 – Cell typing of a frozen kidney tumor, related to Figure 3**

**Figure S5 – Cell type specific genes with DCIs, related to Figure 4**

**Figure S6 – Shared mutation hotspots in all major cell types from a frozen kidney tumor, related to Figure 5**

**Figure S7 – Single cell transcriptome-wide mutations in a frozen kidney tumor. Related to Figure 5.**

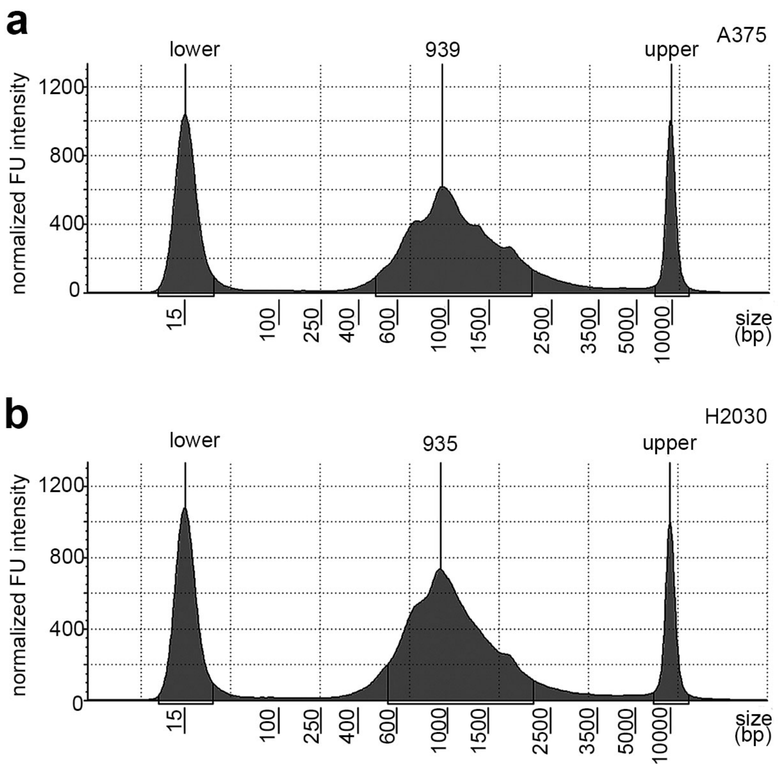


**Figure S1 – Size distributions of cDNAs, related to Figure 1-2**
The TapeStation traces of full-length cDNAs of **a**, A375 and **b**, H2030 before making sequencing libraries.

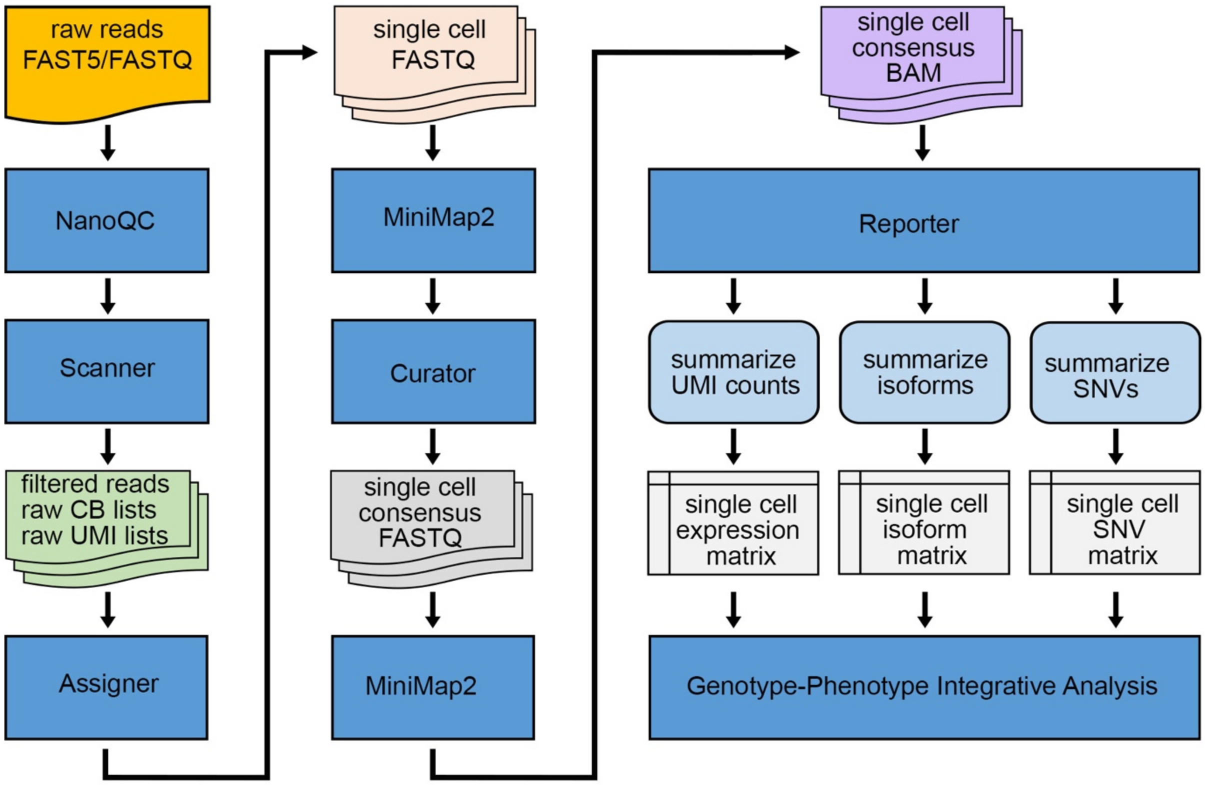


**Figure S2 – Illustration of scNanoGPS methods, related to Figure 1**
Blue boxes represent execution items, while others indicate files.

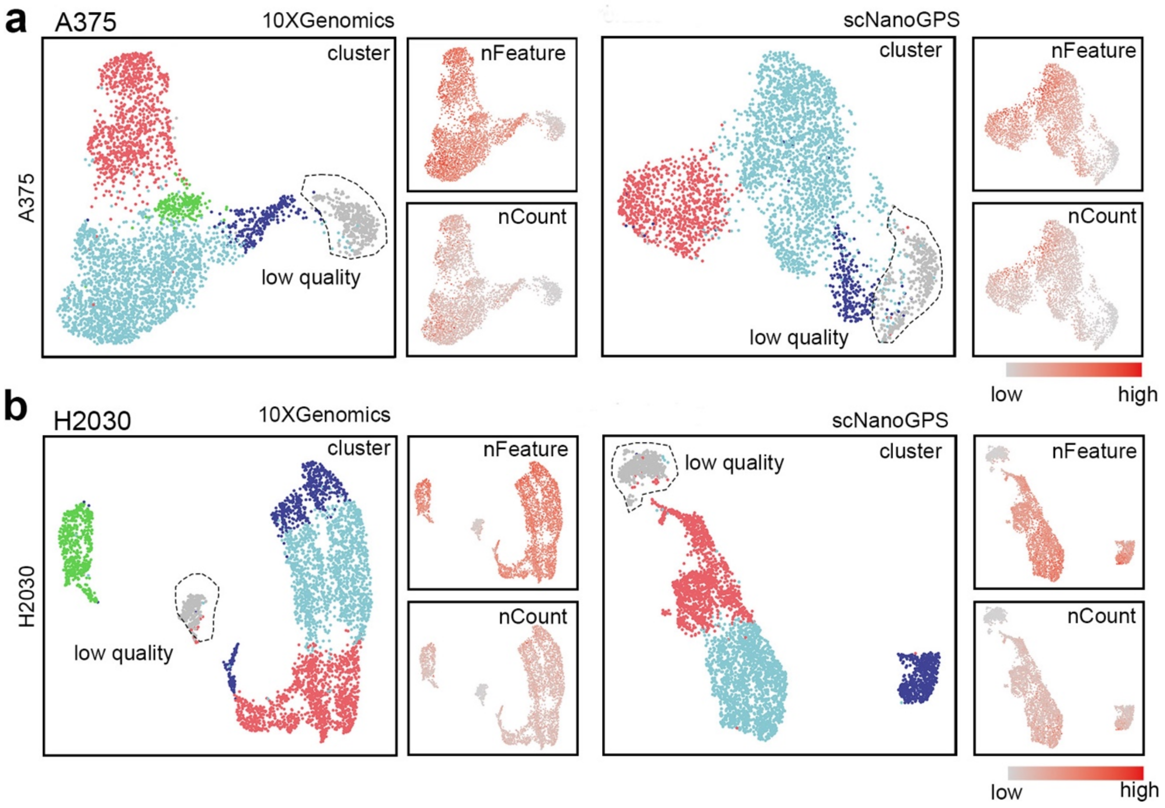


**Figure S3 – Identification of low-quality cells, related to Figure 2**

**a**, UMAPs of low-quality cells detected by NGS and scNanoGPS in A375. **b**, UMAPs of low-quality cells detected by NGS and scNanoGPS in H2030.

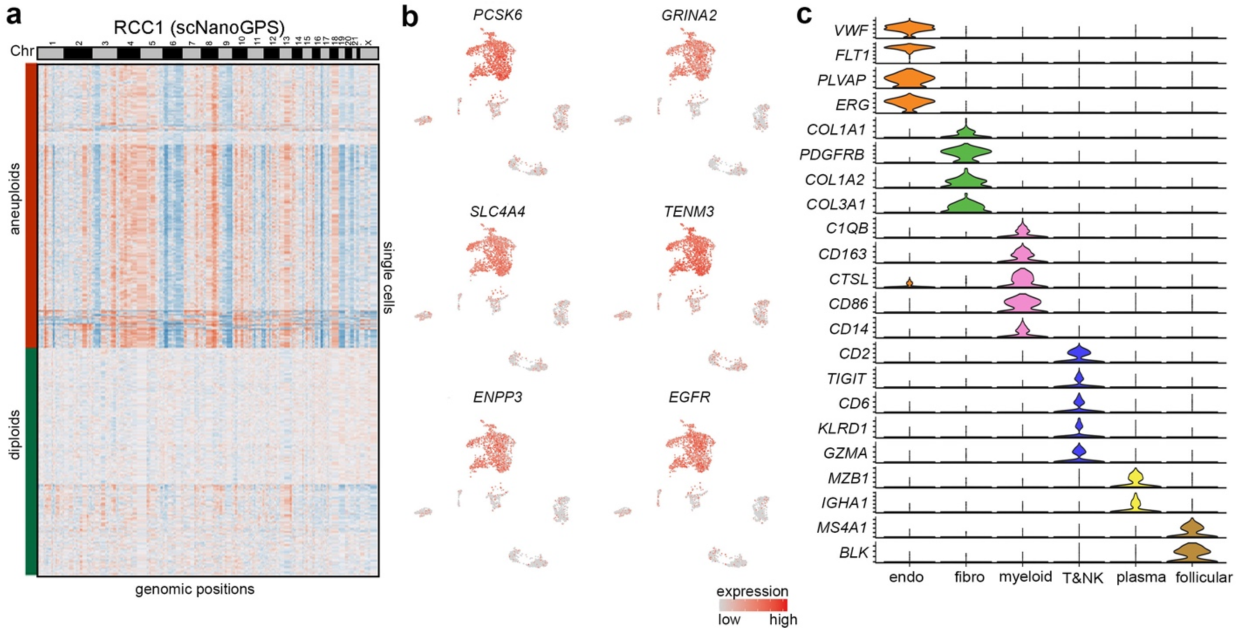


**Figure S4 – Cell typing of a frozen kidney tumor, related to Figure 3**

**a,** Heatmap of single cell copy number profiles with long-read transcriptome data calculated by CopyKAT. **b**, Gene expression UMAPs of known kidney cancer genes. **c**, Violin plots of known cell type-specific marker genes.

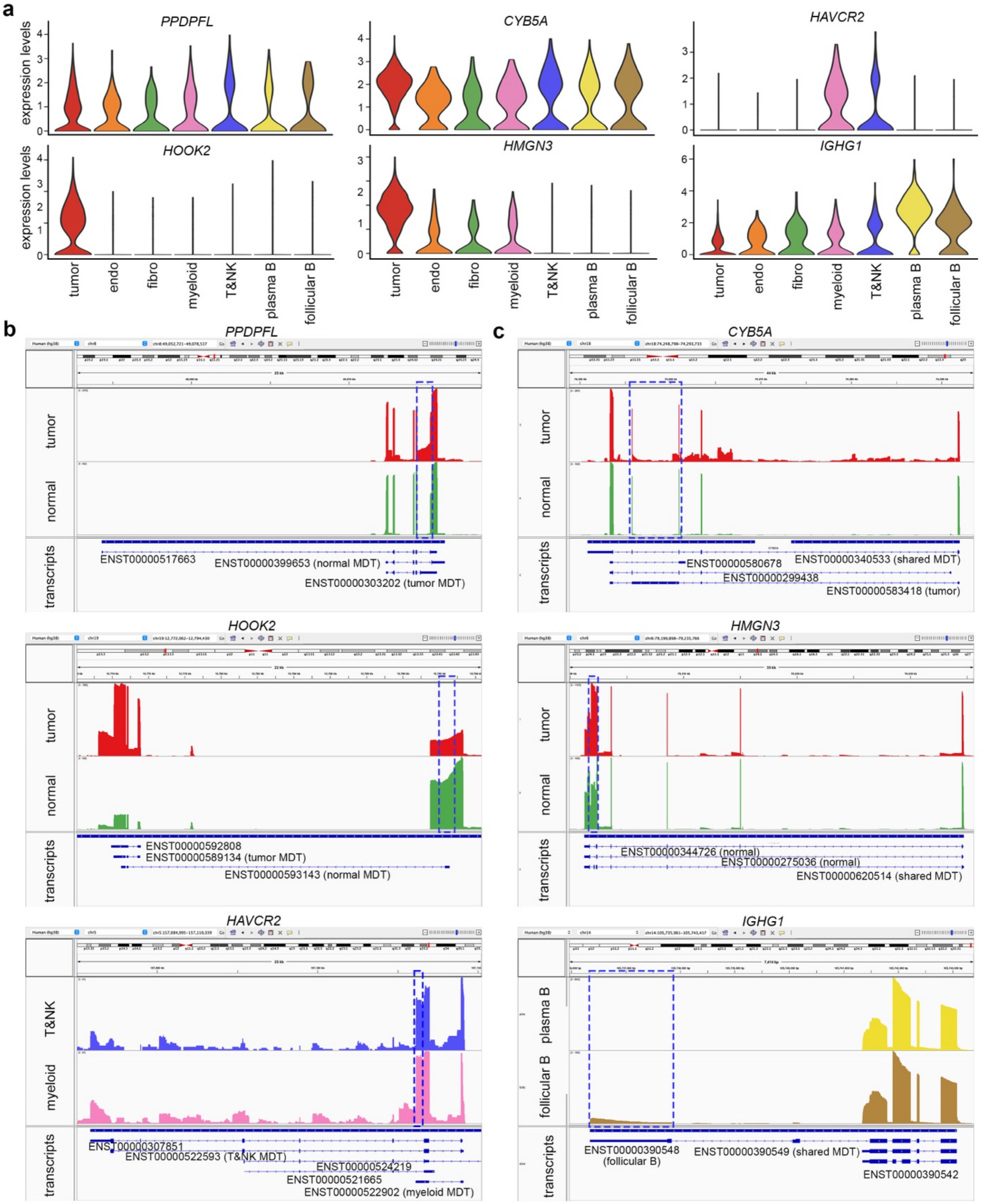


**Figure S5 – Cell type specific genes with DCIs, related to Figure 4**

**a**, Violin plots of gene expression levels of 6 example genes with cell-type-specific DCIs. **b**, IGV visualization of reads mapped to 3 example DCIs genes expressing different MDTs in different cell types. **c**, IGV visualization of reads mapped to 3 example DCIs genes expressing same MDTs in different cell types. Shown reads included both pre-mRNA and mRNA mapping reads.

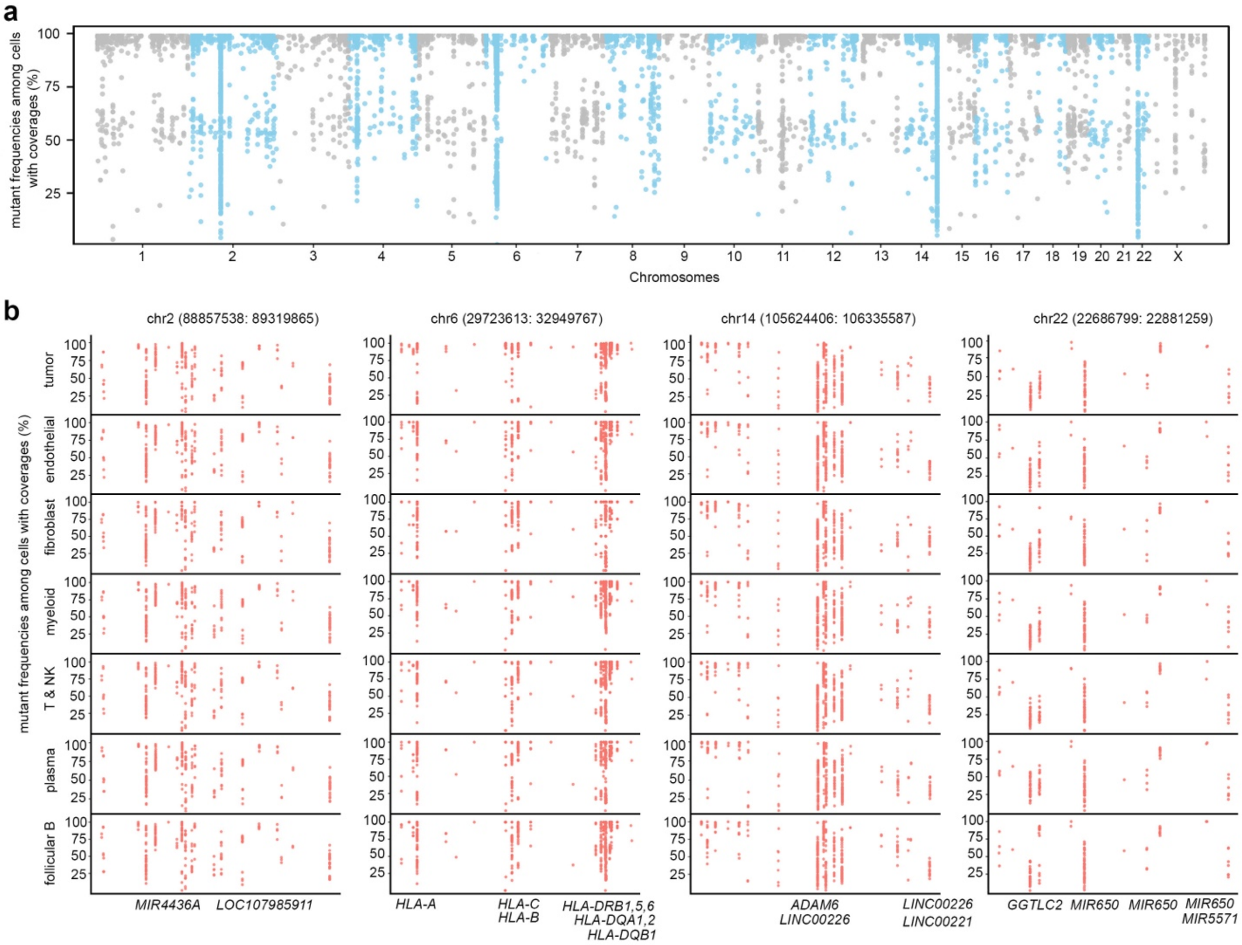


**Figure S6 – Shared mutation hotspots in all major cell types from a frozen kidney tumor, related to Figure 5**
**a**, Cellular frequencies of all mutations, i.e., percentages of cells expressed mutants over all cells that had coverages.
**b**, Cellular frequencies of mutations located in 4 hot spots in each major cell type.

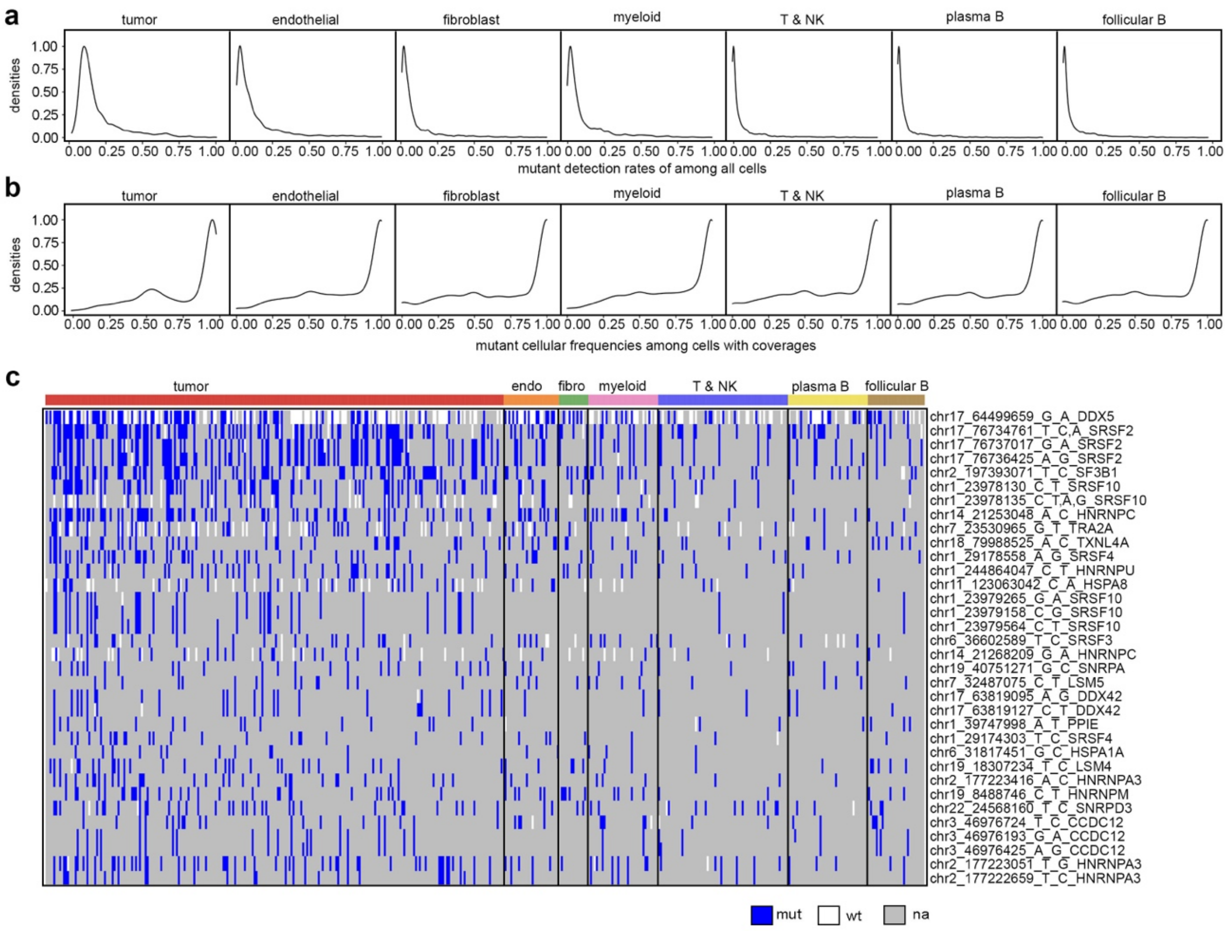


**Figure S7 – Single cell transcriptome-wide mutations in a frozen kidney tumor. Related to Figure 5**
**a**, Overall detection rates of all mutations, i.e., cells expressed mutants over all cells within in each cell type. **b,** Mutation cellular frequencies, i.e., cells expressed mutants over cells with coverages in each cell type. **c**, Heatmap of single cell mutations located in SPLICESOME genes.
